# A multistable slow-fast model of affective state switching under circadian drive

**DOI:** 10.64898/2026.02.11.705337

**Authors:** Veronica Wen-Ting Will, Paola Magioncalda, Matteo Martino, Jihwan Myung

**Affiliations:** Graduate Institute of Mind, Brain and Consciousness (GIMBC), Taipei Medical University, New Taipei City 235, Taiwan; International Master/PhD Program in Medicine (IGPM), College of Medicine, Taipei Medical University, Taipei 110, Taiwan

**Keywords:** bipolar disorder, circadian rhythms, diurnal mood variation, Ornstein-Uhlenbeck perturbations, affective dynamics, multistability, slow-fast system, HPA-axis homeostasis

## Abstract

Rhythmic mood fluctuations are often associated with circadian timing, but it remains unclear how daily rhythms that normally organize physiological affective variation also shape transitions into longer pathological episodes in bipolar disorder. We introduce a phenomenological slow-fast model of affective dynamics inspired by HPA-axis homeostatic feedback. A fast affective coordinate evolves on a multistable landscape whose geometry is shaped by a slow homeostatic variable, producing four stable wells: depression-like, physiological low, physiological high, and mania-like. Circadian input acts not as a local forcing term but as a periodic tilt of the global affective landscape. With intact barriers, this tilt entrains physiological low-high cycling; when outer barriers are weakened, it creates phase-dependent windows for stochastic escape into pathological wells, where prolonged occupancy can produce episode-like infradian cycling. Modeling persistent perturbations as an Ornstein-Uhlenbeck process, we derive a Kramers-type escape criterion linking pathological transitions to phase-dependent barrier height and perturbation strength. One-year simulations and scans over the parameter governing landscape geometry show that preserved circadian amplitude stabilizes physiological cycling, whereas reduced amplitude combined with stronger perturbations promotes entry into and occupancy within depression-like or mania-like wells. The model provides a dynamical framework for studying how circadian disruption, landscape geometry, and perturbation sensitivity jointly shape transitions between physiological mood cycling and episode-like pathological states.

**Lead Paragraph:** Bipolar disorder involves large, persistent transitions among depressive, euthymic, and manic states, whereas everyday mood also shows smaller fluctuations over the day. Whether these two forms of affective variation can be understood within a single dynamical framework remains unclear. Here, we present a slow-fast affective landscape model in which a slow homeostatic variable shapes a multistable potential for a faster affective state. The landscape contains two inner physiological wells and two outer pathological wells. Circadian input does not force the affective state directly into particular wells; instead, it periodically tilts the global landscape, entraining daily cycling within the physiological low and high states under normal geometry. When circadian amplitude is weakened or outer barriers are reduced, persistent stochastic fluctuations can promote escape into hard-to-exit depression-like or mania-like states. The central issue is therefore not simply whether such escapes can occur, but how circadian disruption, landscape geometry, and perturbation sensitivity determine the balance of dwell time across physiological and pathological states over clinically meaningful observation windows.

## 1. INTRODUCTION

Bipolar disorder encompasses severe, recurrent, and often chronic psychiatric conditions characterized by transitions among depressive, euthymic, hypomanic, and manic states (American Psychiatric Association, 2022; Grande et al., 2016; Carvalho et al., 2020; McIntyre et al., 2020). In DSM-5-TR, bipolar disorder type I (BD-I) is diagnostically anchored by at least one manic episode, whereas major depressive episodes are common but not required for the diagnosis (American Psychiatric Association, 2022). In prototypical presentations, mania features euphoric or irritable mood and increased activity or energy, whereas depression features depressed mood and/or anhedonia, placing these states at opposite poles of affective pathology. Clinically, however, manic and depressive episodes are distinguished from ordinary mood fluctuation by their severity, persistence, symptom burden, and functional consequences.

An important clinical asymmetry is that the diagnostic salience of mania does not imply that manic states dominate the longitudinal course of illness. Even in BD-I, depressive symptoms often account for more time and morbidity than manic or hypomanic symptoms (Judd et al., 2002; Grande et al., 2016; McIntyre et al., 2020). This distinction is central for dynamical modeling. A model of bipolar affective dynamics should not merely represent two symmetric poles of mood. It should account for regular physiological affective variation, rare but clinically consequential transitions into mania-like states, and potentially larger cumulative occupancy in depression-like states. In this sense, bipolar disorder motivates a dwell-time formulation: the clinically relevant question is not only whether pathological states are reachable, but how long the system remains in physiological versus pathological regions over meaningful observation windows.

Mathematical models of bipolar disorder have used limit cycles, coupled oscillators, autoregressive processes, and bistable switching frameworks (Daugherty et al., 2009; Goldbeter, 2011; Steinacher and Wright, 2013; Nunes et al., 2022). These models formalize recurrence, oscillation, and state switching, but many remain largely abstract. Conversely, biologically grounded feedback models, including endocrine models, provide mechanistic structure but do not directly encode the clinically observed coexistence of diurnal mood variation and episode-like affective extremes. This motivates an intermediate approach: a biologically inspired slow-fast system in which the fast state variable is interpreted as an affective coordinate rather than as a directly measured hormone concentration.

Circadian regulation provides a natural organizing input for such a model. Circadian rhythm disturbances are highly prevalent in individuals with bipolar disorder and have been linked to sleep-wake timing, light sensitivity, seasonality, endocrine regulation, and episode vulnerability (Takaesu, 2018; Walker et al., 2020; Dollish et al., 2024; Rahim et al., 2025). Diurnal variation in mood symptoms is central in specific clinical subtypes; notably, melancholic depression is characterized by marked circadian abnormalities, including a consistent worsening of symptoms in the early morning hours (American Psychiatric Association, 2022). Abrupt circadian misalignment after transmeridian travel has been associated with polarity-specific mood changes (Inder et al., 2016; Walker et al., 2020; Rahim et al., 2025). Chronic circadian disruption by night-shift work alters sleep-wake and endocrine timing, and a meta-analysis of observational studies found that night-shift workers had an approximately 40% higher risk of depression than non-night-shift workers (Lee et al., 2017; Walker et al., 2020). These observations motivate a dynamical framework in which the amplitude, phase, and regularity of circadian input can reshape affective-state occupancy, from physiological circadian-scale variation to pathological infradian-scale persistence in bipolar disorder.

Hypothalamic-pituitary-adrenal (HPA)-axis regulation provides one biological formulation for this slow-fast system. Cortisol and other endocrine signals exhibit strong diurnal variation under suprachiasmatic nucleus control, and HPA-axis dysregulation has repeatedly been associated with affective disorders (Adam et al., 2017; Jones et al., 2021; Herane-Vives et al., 2020; Huang et al., 2017; Murri et al., 2016; Milo et al., 2024). At the same time, endocrine findings in bipolar disorder are heterogeneous across episode polarity, sampling modality, medication status, and clinical context (Stetler and Miller, 2011; Murri et al., 2016; Maripuu et al., 2014). These mixed findings argue against a direct one-to-one identification of cortisol, corticotropin-releasing hormone (CRH), or other endocrine variables with mood state. Accordingly, we use HPA-axis homeostatic feedback to motivate the model architecture and interpret the modeled fast variable as a phenomenological affective coordinate.

Motivated by this clinical and circadian background, we introduce a quadristable slow-fast model of affective-state switching under circadian drive. The model has two inner wells for physiological low and physiological high states and two outer wells for depression-like and mania-like pathological states. Circadian input does not act as a local force that pushes mood toward a prescribed state. Instead, it periodically tilts the multistable affective potential, so barrier heights and transition accessibility change over the circadian cycle. Ornstein-Uhlenbeck (OU) noise represents a persistent, mean-reverting stochastic load on the fast affective coordinate rather than independent instantaneous impulses. Biologically, it is interpreted as stressor load, sleep disruption, social-rhythm disruption, or arousal-related carryover with finite temporal autocorrelation (see Sec. 2.3 for formal definition and rationale).

The central analytical contribution is to separate three processes that are often treated together: physiological entrainment, pathological entry, and pathological state occupancy. Physiological regulation requires circadian-scale switching between the two inner wells, whereas pathological episodes correspond to rare escapes into the outer wells followed by prolonged occupancy. We therefore pair one-year fractional occupancy simulations with a cycle-integrated, Kramers-type escape criterion linking pathological transitions to phase-dependent barrier height and perturbation strength. This formulation also clarifies the relation to stochastic resonance: the desirable process is physiological circadian-scale switching, whereas pathological escape should remain slower than the circadian frequency. Throughout, “mania-prone,” “euthymia-like,” and “BD-I-like” denote model-occupancy regimes rather than clinical classifications. The BD-I-like regime is one in which mania-like pathological-high (PH) state occupancy remains rare but reachable, depression-like pathological-low (PL) state occupancy carries the larger share of pathological burden, and physiological state occupancy remains substantial. This dwell-time formulation reframes the clinical question: not whether pathological states can be reached in principle, but which factors sustains physiological state occupancy over clinically meaningful observation windows. In the simulated regimes, reduced circadian amplitude together with stronger stochastic load shifts affective dynamics from circadian-scale physiological variation toward infradian, depression-weighted pathological dwell time resembling the longitudinal burden observed in BD-I.

## 2. MODEL FORMULATION

### 2.1. Reduced slow-fast model

Our model describes affective dynamics on two timescales, represented by a slow homeostatic variable and a fast affective variable. This slow-fast framework is well suited to biological systems in which rapid dynamics are modulated by slower regulatory mechanisms. We begin from nondimensional HPA-axis models in which stored CRH, circulating CRH, ACTH, cortisol, and glucocorticoid receptors interact through feedback loops capable of bistability (Jelić et al., 2005; Gupta et al., 2007; Andersen et al., 2013; Kim et al., 2016; Cheng et al., 2021). Following the reduction in Cheng et al. (2021), the downstream pituitary-adrenal variables are placed at quasi-equilibrium for a given circulating CRH level. This yields a two-variable slow-fast feedback architecture. In the present model, the rescaled slow variable *x* represents homeostatic or endocrine load, and the fast variable *y* represents affective state.

The slow equation is

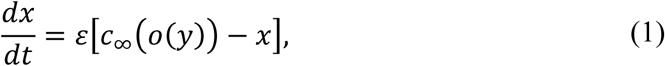

where *c*_min_ is the constant minimal stored-CRH level and *c*_∞_(*o*) is the input-dependent slow-nullcline function:

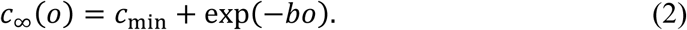

The cortisol-related proxy *o*(*y*) is not integrated as an independent state variable. It is obtained from a quartic constraint inherited from the quasi-steady-state reduction. We assume that *a*, *o*, and *r* are at equilibrium within this reduced system and obtain *o*(*y*) by setting each of their respective derivatives to zero. Writing *q* = *o*^2^, the constraint is quadratic in *q*:

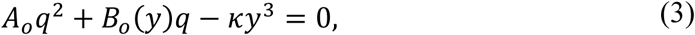

with

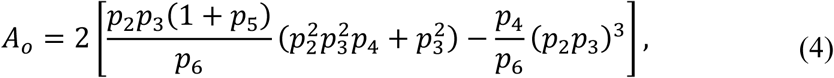

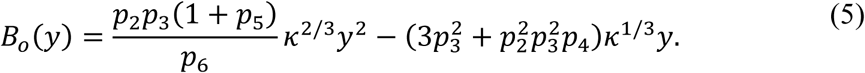

The positive real solution is then

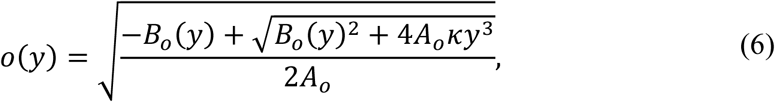

when the radicand and numerator are positive. This form makes clear why the cortisol-related proxy is quartic in *o*: it is the algebraic consequence of eliminating ACTH, cortisol, and receptor variables from the reduced HPA-axis formulation, not an additional empirical fit imposed on mood.

The quasi-steady reduction is a modeling approximation. It assumes that downstream endocrine variables relax quickly relative to the retained slow homeostatic coordinate. We therefore use this reduction only to motivate the geometry of the affective landscape; after the multistable extension, *x*, *y*, and *o*(*y*) are not interpreted as directly measured hormone concentrations.

### 2.2. Quadristable affective nullcline

To represent both physiological and pathological affective states, we expand the fast nullcline from a cubic bistable form to a septic polynomial. Let *a*_1_,…, *a*_6_ denote six prescribed turning-point locations of the fast nullcline and let *S_m_* be the elementary symmetric sum of order *m* over these turning points, with *S*_0_ = 1. These *a_i_* are roots of *dP*_7_/*dy*; they are not equilibria of the full slow-fast system. Define

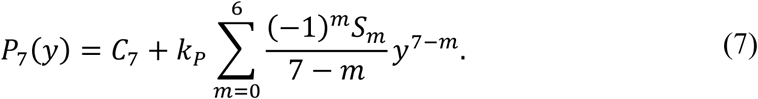

Equivalently,

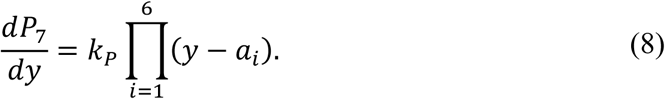

The integration constant *C*_7_ is part of the nullcline definition because it vertically shifts the fast nullcline and can change the number and location of intersections. All simulations use the nondimensional convention *C*_7_ = 0 and *k_P_* = 1, so *P*_7_(0) = 0. The inherited HPA scaling enters through the fast-vector-field gain *γ* = *κ*^−1/3^ rather than through an additional hidden nullcline offset. The deterministic fast equation is

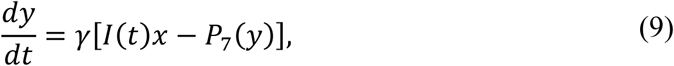

so the fast nullcline is *I*(*t*)*x* = *P*_7_(*y*) and is unchanged by the positive gain *γ*.

The seventh-order form is phenomenological, but it is not arbitrary. In a one-dimensional potential, four stable wells require four minima separated by three intervening local maxima, therefore requiring that the fast equation produces seven fixed points along *y* for a fixed value of the slow coordinate. A seventh-order nullcline is the minimum-degree polynomial construction that can create those seven intersections with alternating stability. Although higher-order polynomials could generate additional wells, we use the seventh-order model to consider four affective states: pathological low (PL), physiological low (L), physiological high (H), and pathological high (PH). The construction is intentionally minimal. Other numbers of stable states could be constructed in principle, and the model does not claim that quadristability is uniquely determined by the underlying biology. The four wells are chosen to encode the clinical distinction between physiological diurnal variation and two pathological poles.

The parameter *a*_3_ is one of the turning points of the septic fast nullcline, i.e., varying *a*_3_ bends the middle fold and rearranges the spacing between the physiological attractors and the nearby saddles. This changes the relative accessibility of PL and PH, so *a*_3_ acts as a geometric central-bend parameter (**Fig. 1A**). Circadian input, by contrast, periodically shifts the fast nullcline relative to the slow nullcline and changes the instantaneous flow and basin accessibility at a given circadian phase (**Fig. 1B**). This is equivalent to tilting the potential landscape, which biases overall episode polarity and gates phase-dependent escape risk (**Fig. 1C**).

**Figure 1.**
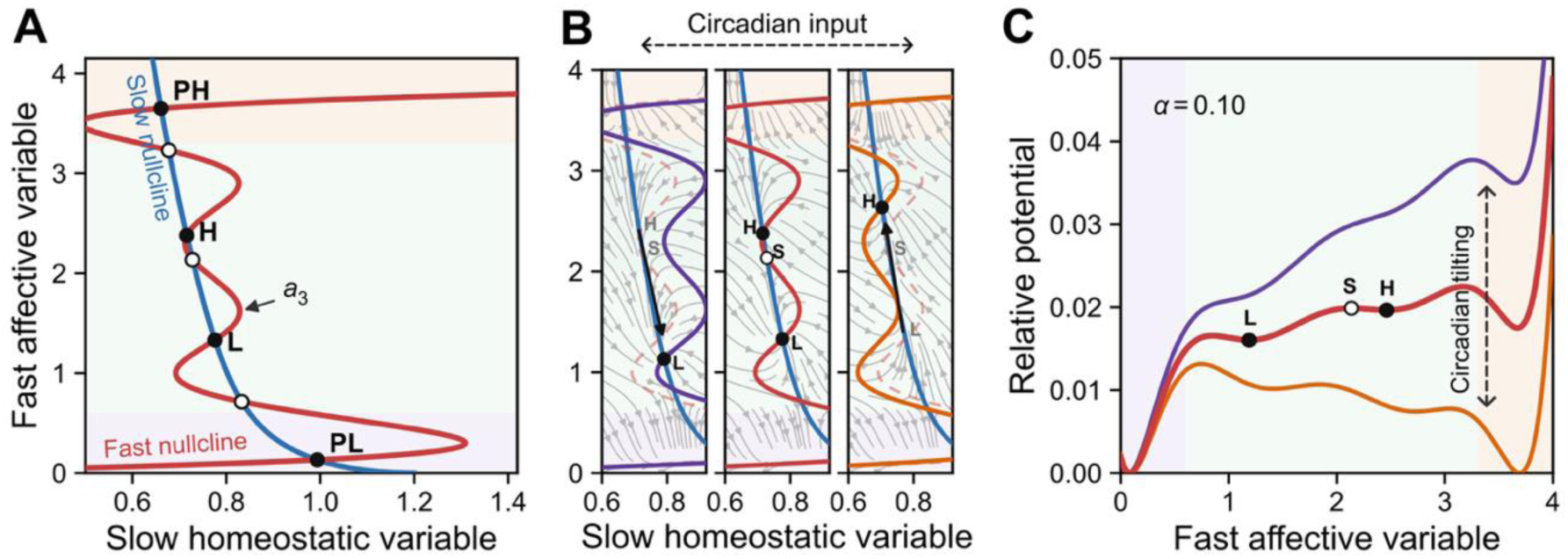
Baseline geometry, phase flow, and circadian tilting of a four-state affective landscape. (**A**) Phase-plane geometry of the deterministic slow-fast system. The solid red curve is the fast nullcline, and the blue curve is the slow nullcline. Their intersections produce four stable fixed points (filled circles) that alternate with three saddle points (open circles). The four stable states lie within the pathological-low (PL), physiological-low (L), physiological-high (H), and pathological-high (PH) affective regions. The parameter *a*_3_ marks the central bend of the fast nullcline near the physiological wells. (**B**) Fixed-phase deterministic flow fields for trough (left), midline (middle), and peak (right) circadian input, with OU perturbation set to zero. Gray streamlines indicate instantaneous deterministic drift. Circadian input shifts the fast nullcline relative to the slow nullcline, moving the stable points (L, H) and saddle points (S) at each circadian phase. (**C**) Relative fast-potential profiles at the fixed slow coordinate *x* ≈ 0.738. The black curve is the baseline relative potential at *I* = *I*_0_; filled circles mark its local minima, and open circles mark the intervening barriers. The translucent envelope shows the phase-dependent tilt over one circadian cycle for a small illustrative amplitude, *I*(*φ*) = *I*_0_ + *α* sin*φ*, with *α* = 0.10. Shaded regions follow the PL, physiological, and PH convention in (A).

### 2.3. Circadian drive and OU perturbations

Circadian input is introduced as

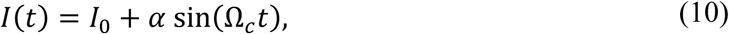

here *α* is the circadian amplitude and *φ*(*t*) = Ω*_c_t* is the circadian phase. Unless otherwise stated, time is reported in physical units, with *T_c_* = 1 day such that Ω*_c_* = 2*π*/*T_c_*. Numerical integrations are performed in model time and then rescaled to physical time so that one full cycle of *I*(*t*) corresponds to one day (150.8 model-time units). The one-year simulations contain 365 circadian cycles.

Because *I*(*t*) is defined as a positive input, the main parameter scans are restricted to 0 ≤ *α* ≤ *I*_0_. With *I*_0_ = 1, this corresponds to 0 ≤ *α* ≤ 1. Values *α* > *I*_0_ would make the input negative during the trough and are not included in the main biologically interpretable scan. Circadian input through *I*(*t*) causes the system to stably oscillate between the two physiological affective wells (**Fig. 2A,B**).

**Figure 2.**
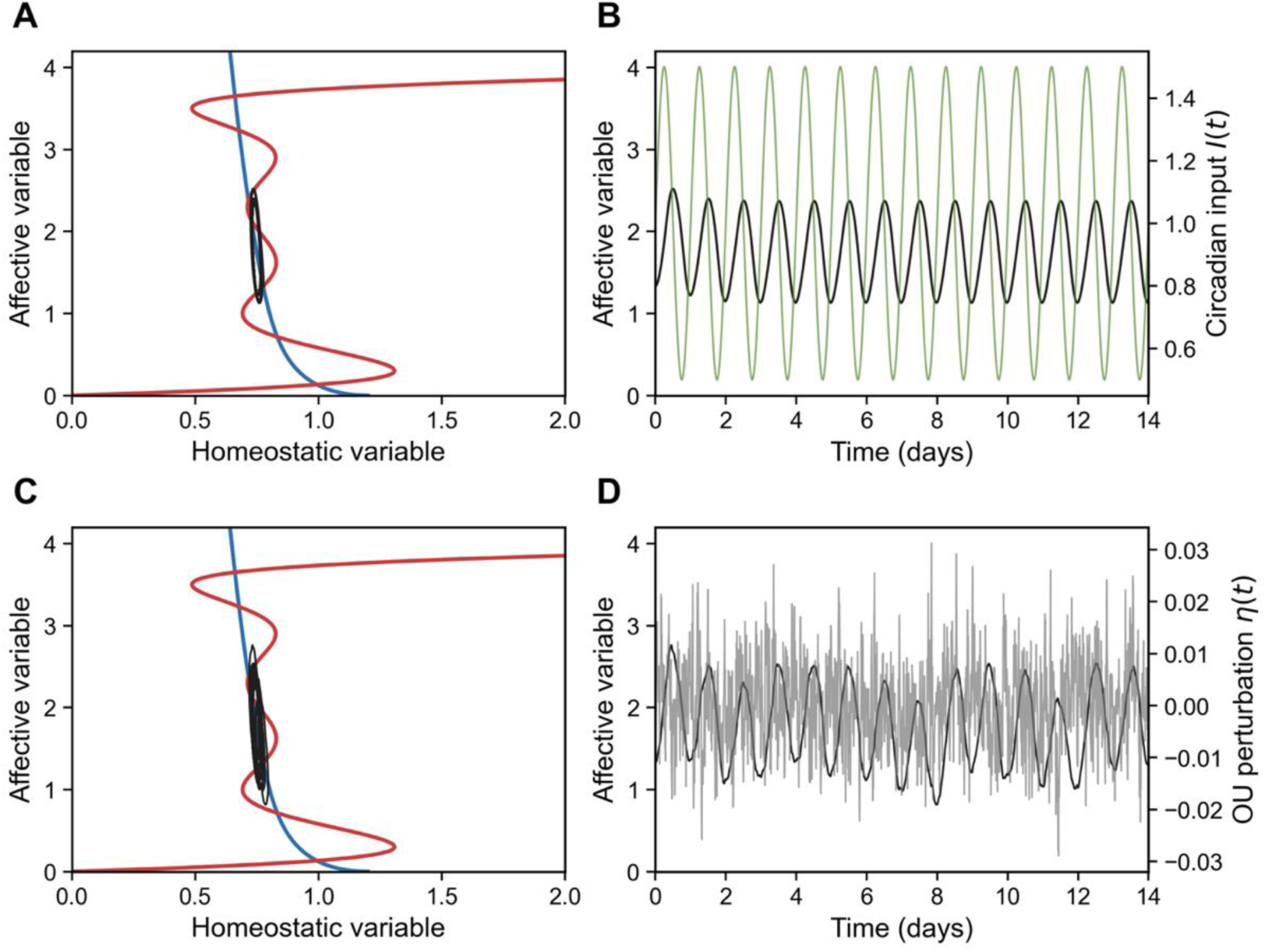
Circadian confinement of the physiological affective pair with and without OU perturbations. (**A,B**) Stable affective state alternations between physiological low and high states, entrained to circadian driving at preserved amplitude (*α* = 0.5) in the absence of perturbations. The left panel shows the phase-plane trajectory on the slow homeostatic and fast affective axes (**A**), and the right panel shows the corresponding fast affective coordinate over time (**B**). (**C,D**) Adding low-intensity zero-mean OU perturbations (*σ_η_* = 0.006, *θ* = 0.3) produces visible fluctuations around the same circadian path but no sustained occupancy in the outer pathological wells over the simulated interval.

Persistent perturbations are modeled with a zero-mean OU process on the fast coordinate, where *θ* is the mean-reversion rate and *σ_η_* is the perturbation intensity:

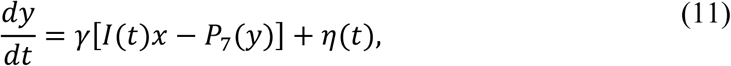

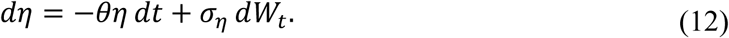

The OU term is used because stress, arousal, sleep loss, social stimulation, and circadian misalignment usually persist for finite durations and decay gradually toward baseline. They are therefore not well represented as independent impulses at every instant. The finite memory of the OU process means finite temporal autocorrelation. Its stationary autocovariance is

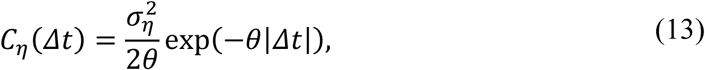

so the characteristic perturbation persistence time is *τ*_OU_ = *θ*^−1^. Because the OU drift returns *η* to zero, the stochastic perturbation does not by itself generate a sustained manic-like (PH) or depressive-like (PL) bias when perturbation intensity is low. Polarity bias is instead introduced geometrically through the nullcline-bend parameter *a*_3_. Adding low OU perturbation (*σ_η_* = 0.006) produces minor random fluctuations within the physiological affective wells over time (**Figs. 2C,D**).

### 2.4. Effective potential and escape criterion

For a fixed slow coordinate *x* and circadian phase *φ*, the fast stochastic equation can be written as motion in an effective one-dimensional potential:

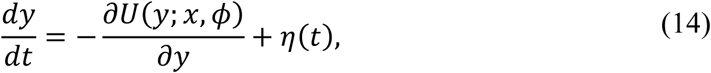

where

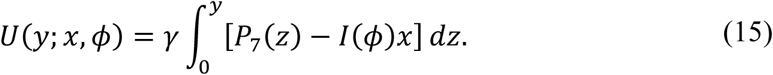

The lower integration limit in Eq. (15) fixes only the arbitrary zero of *U*. For visualization, all potential plots use the phase-specific relative potential *U*_rel_(*y*; *x*, *φ*) = *U*(*y*; *x*, *φ*) − min*_y_U*(*y*; *x*, *φ*) where the minimum is taken over the plotted fast-coordinate range for the same fixed *x* and *φ*. This normalization sets the lowest point of each profile to zero and allows comparison on a common vertical scale, while fixed points and within-profile barrier heights remain untouched since *∂_y_U*_rel_ = *∂_y_U*. Escape barriers are therefore interpreted as differences between a well minimum and an adjacent saddle within the same circadian phase. Circadian forcing acts through the linear term −*γI*(*φ*)*xy*, producing a periodic tilt of the affective landscape. For modest tilts, it primarily changes barrier heights; for larger instantaneous tilts, it can also remove or restore local wells in a potential cross-section (**Fig. 1**). This phase-dependent reshaping is the gating mechanism tested by direct simulation.

For an illustrative system with *a*_3_ = 1.593, the potential is analyzed at the center saddle where *x* ≈ 0.738 (**Fig. 3**). The potential is also analyzed at the two physiological stable points, where *x* ≈ 0.708 for the H state and *x* ≈ 0.773 for the L state. In this cross-section, the landscape creates a large stable attractor with two smaller pockets of stability within it. From the middle saddle, the system can move in either direction with equal energetic favorability. Once the system reaches a physiological high or low stable point, the most favorable direction of travel is back toward the center. The relatively low potential increase needed to leave one circadian state and move to the other means that a small change in circadian input *I*(*t*) provides the potential change required for state switching. Circadian forcing therefore acts as a periodic tilt of the affective landscape, changing barrier heights without changing the underlying nullcline topology.

**Figure 3.**
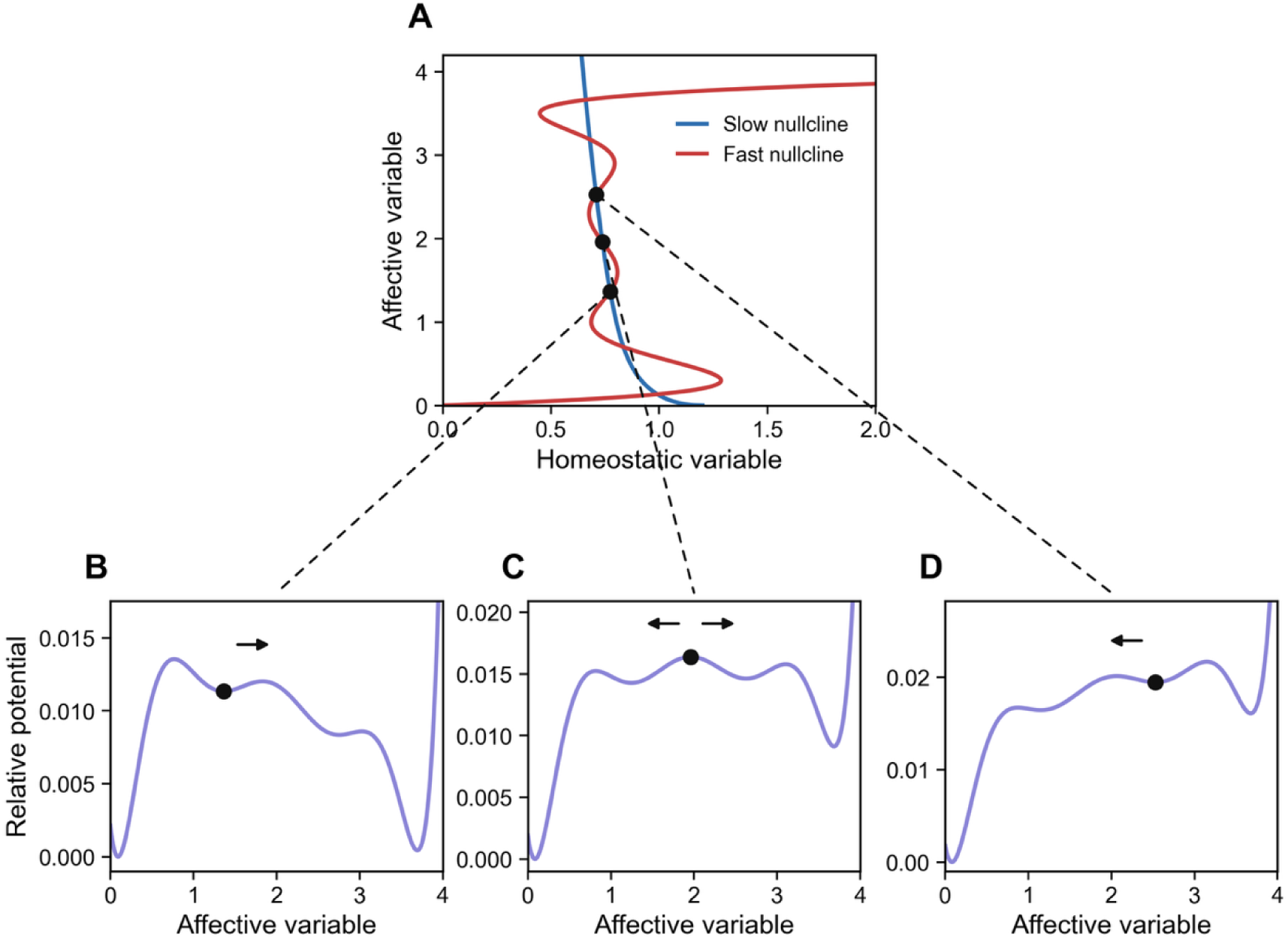
Potential landscape interpretation of the three middle intersections of the nullclines. (**A**) Phase plane showing nullclines of the slow homeostatic variable (blue) and fast affective variable (red). Filled circles mark fixed points. (**B**-**D**) Relative potentials on the nullclines for the three fixed *x*-values in the absence of circadian drive. At the middle saddle (*x* ≈ 0.738), the intersection is unstable, and the system can travel either up or down with equal favorability, corresponding to equal probability of motion to either neighboring well (**C**). For the intersections at *x* ≈ 0.708 (physiological high; **B**), and *x* ≈ 0.773 (physiological low; **D**), returning toward the other physiological stable state is much more favorable than escaping to a pathological state.

Let *y_p_*(*t*) be the occupied physiological well *p* and *y_s_*(*t*) the adjacent saddle *s* separating this well from an outer pathological well. The phase-dependent pathological escape barrier to the destination well *d* is

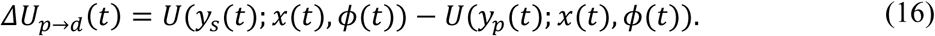

In the weak-noise, rare-escape regime, a local Kramers-type rate approximation summarizes the fixed-phase escape tendency,

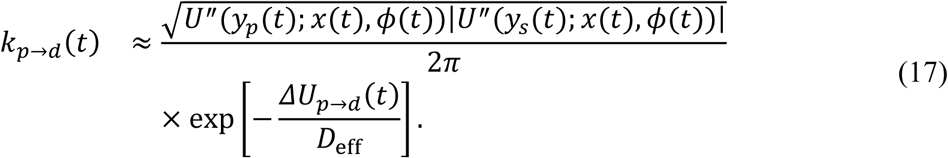

We use this expression as a hazard proxy rather than as a substitute for the direct stochastic simulations. Specifically, the exponential factor depends on the ratio of the escape barrier height *ΔU_p_*_→*d*_(*t*) to the effective noise scale *D*_eff_. For *ΔU_p_*_→*d*_(*t*) ≫ *D*_eff_, escape becomes rare. The square-root prefactor accounts for the local curvature at the physiological minimum and the adjacent saddle; that is, it reflects how tightly the well confines the trajectory. When the OU correlation time is short relative to the barrier-crossing timescale, the effective diffusion scale can be approximated by the integrated autocovariance of the OU process:

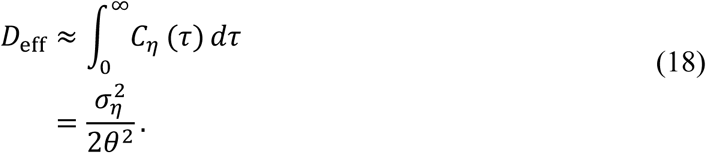

In practice, the escape rate is evaluated separately for the two adjacent pathological exits, from the physiological well to PL (*p* → PL) and to PH (*p* → PH), and the total pathological escape hazard is their sum when both exits are accessible. When OU persistence is comparable to the circadian or barrier-crossing timescale, Eqs. (17)-(18) should be read as a phase-resolved escape-rate approximation. Direct stochastic simulations are therefore the primary dwell-time calculation. The expected number of pathological escapes during one circadian cycle is the integrated hazard

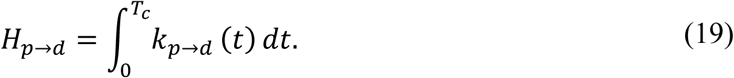

Physiological confinement requires *H_p_*_→*d*_ ≪ 1. Because the simulations span many circadian cycles, the per-cycle hazard should not be interpreted alone. Under the rare-event approximation, if the simulated window contains *N* circadian cycles, then the probability of at least one pathological escape over that window is

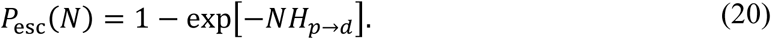

Thus *H_p_*_→*d*_ ≪ 1 indicates rare escape per cycle, whereas sustained confinement over the full simulated window requires *NH_p_*_→*d*_ ≪ 1. Regular physiological alternation requires a different condition: transition time between the two inner wells should be comparable to a circadian half-cycle, whereas pathological escape should remain much slower than the circadian cycle. The analytical stability window is therefore selective physiological switching with rare outer-well entry.

### 2.5. Fractional occupancy calculations

Because the absence of pathological escape does not necessarily imply healthy dynamics, we quantify fractional occupancy across the four affective wells over one-year simulations. Time points are classified by fixed affective-coordinate thresholds corresponding to the shaded regions in **Figs. 1** and **4**-**8**: PL, *y* < 0.60; L, 0.60 ≤ *y* < 2.0; H, 2.0 ≤ *y* ≤ 3.30; and PH, *y* > 3.30. Healthy physiological cycling corresponds to occupancy distributed across the two physiological wells with little outer-well dwell time. Static low-amplitude regimes can show little pathological dwell time but also little physiological alternation. Candidate pathological regimes show increased dwell time in one or both outer wells. The parameter scans therefore report the directly simulated fractional occupancies *p*_PL_, *p*_L_, *p*_H_, and *p*_PH_ rather than an additional two-state episode-burden formula.

**Figure 4.**
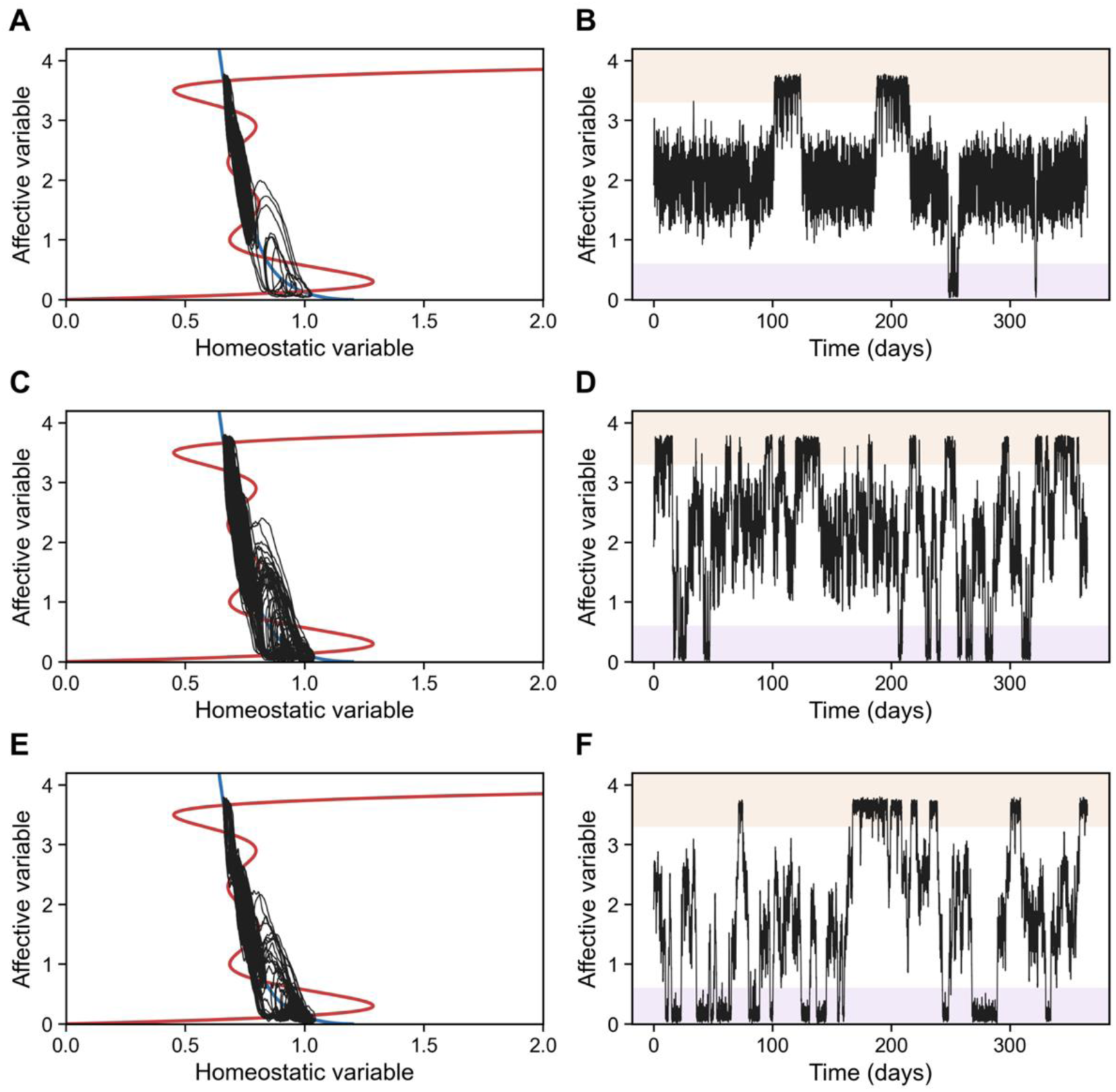
Representative one-year transitions in a candidate mania-prone landscape. Simulations use the mania-prone contrast geometry (*a*_3_ = 1.595, *θ* = 0.3), and each panel shows a single representative stochastic realization. The left column (**A,C,E**) shows the phase-plane trajectory over the deterministic nullclines, and the right column (**B,D,F**) shows the corresponding fast affective coordinate over 365 days. In the time series, shaded bands mark the PL and PH regions. (**A,B**) Preserved circadian amplitude (*α* = 0.5) with low OU perturbation intensity (*σ_η_* = 0.006) keeps state occupancy mostly physiological, with occasional PH visits. (**C,D**) Preserved amplitude (*α* = 0.5) with stronger perturbation (*σ_η_* = 0.012) raises PH access but still permits repeated return to the physiological region. (**E,F**) Reduced circadian amplitude (*α* = 0.3) with the same stronger perturbation further raises occupancy in the PH well. The OU perturbation has zero mean; the bias toward the high pole arises from the landscape geometry set by *a*_3_.

Stochastic simulations used a uniform model-time grid. The deterministic drift was integrated with a fourth-order Runge-Kutta step, and the OU process was updated by Euler-Maruyama with zero mean. Trajectory examples in **Figs. 4**-**6** use Δ*t*_model_ = 1.0, corresponding to 0.00663 days, and start from (*x*_0_, *y*_0_) = (0.738,1.94). Parameter scans in **Fig. 7** use Δ*t*_model_ = 2.0, corresponding to 0.01326 days, and start from (*x*_0_, *y*_0_) = (0.777,1.334). Both initial conditions lie within the physiological region; the difference reflects the representative trajectory starting point versus the scan initialization point, and fractional occupancy was computed over the full one-year window without discarding a burn-in. A proposed step producing nonfinite values or *y* < 0 was rejected, and the previous state was retained for that step; trajectories were not clipped to a lower value or made absorbing at the boundary. This rejection rule is a numerical regularization at the lower boundary, not a model assumption about PL dwell time. **Figs. 4**-**6** show single representative stochastic realizations, whereas **Fig. 7** reports seed-averaged fractional occupancy (8 independent realizations for **Fig. 7B** and 30 independent realizations for **Fig. 7C**).

## 3. RESULTS

### 3.1. Quadristability without circadian drive

With *α* = 0 and the parameter set in Table 1, the fast and slow nullclines intersect seven times, producing four stable fixed points separated by three saddles (S) (**Fig. 1A**). The two inner stable states are interpreted as physiological L and H affective states, whereas the outer stable states are interpreted as depression-like and mania-like states. This interpretation follows from the imposed affective landscape geometry, not from direct identification of CRH or cortisol values with mood. The seventh-order nullcline provides a minimal odd-degree polynomial construction capable of representing four stable affective wells separated by unstable barriers.

**Table 1.**
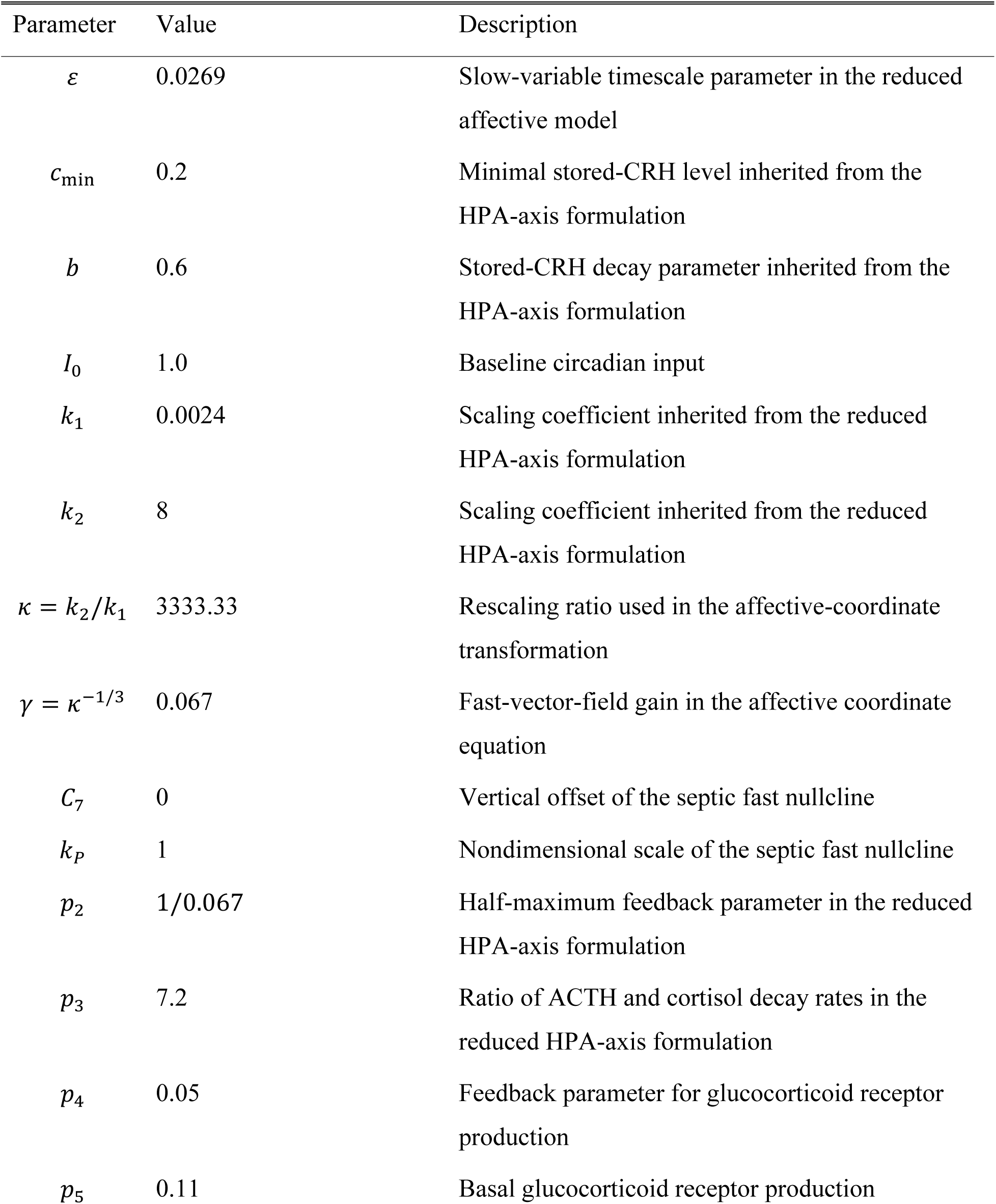
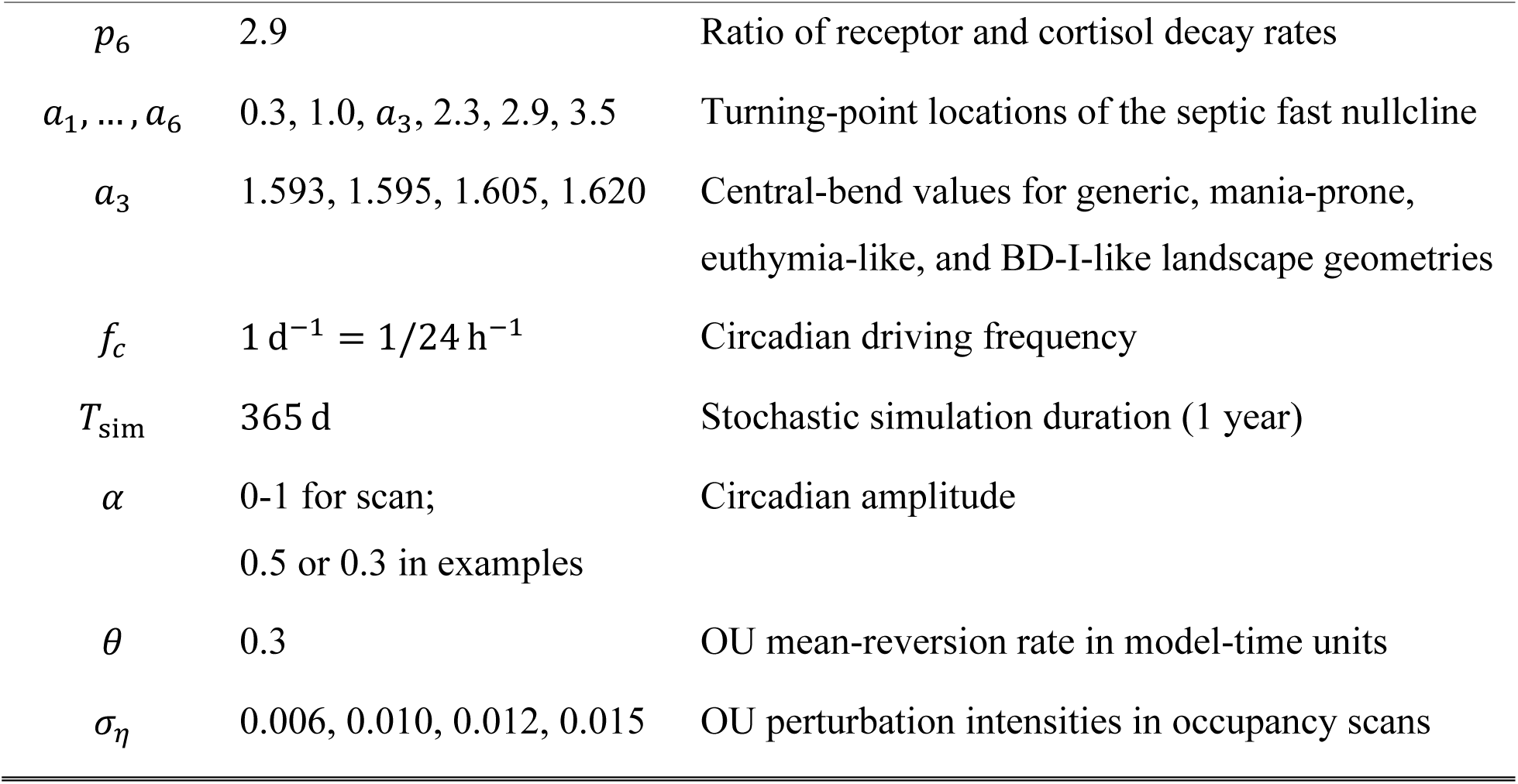
Representative nondimensional parameter values used in the simulations. Parameters are nondimensional unless units are shown; time-dependent results are reported in physical days.

| Parameter | Value | Description |
| --- | --- | --- |
| $\varepsilon$ | 0.0269 | Slow-variable timescale parameter in the reduced affective model |
| $c_{\min}$ | 0.2 | Minimal stored-CRH level inherited from the HPA-axis formulation |
| $b$ | 0.6 | Stored-CRH decay parameter inherited from the HPA-axis formulation |
| $I_0$ | 1.0 | Baseline circadian input |
| $k_1$ | 0.0024 | Scaling coefficient inherited from the reduced HPA-axis formulation |
| $k_2$ | 8 | Scaling coefficient inherited from the reduced HPA-axis formulation |
| $\kappa = k_2/k_1$ | 3333.33 | Rescaling ratio used in the affective-coordinate transformation |
| $\gamma = \kappa^{-1/3}$ | 0.067 | Fast-vector-field gain in the affective coordinate equation |
| $C_7$ | 0 | Vertical offset of the septic fast nullcline |
| $k_p$ | 1 | Nondimensional scale of the septic fast nullcline |
| $p_2$ | 1/0.067 | Half-maximum feedback parameter in the reduced HPA-axis formulation |
| $p_3$ | 7.2 | Ratio of ACTH and cortisol decay rates in the reduced HPA-axis formulation |
| $p_4$ | 0.05 | Feedback parameter for glucocorticoid receptor production |
| $p_5$ | 0.11 | Basal glucocorticoid receptor production |
| $p_6$ | 2.9 | Ratio of receptor and cortisol decay rates |
| $a_1, \dots, a_6$ | 0.3, 1.0, $a_3$, 2.3, 2.9, 3.5 | Turning-point locations of the septic fast nullcline |
| $a_3$ | 1.593, 1.595, 1.605, 1.620 | Central-bend values for generic, mania-prone, euthymia-like, and BD-I-like landscape geometries |
| $f_c$ | $1 \text{ d}^{-1} = 1/24 \text{ h}^{-1}$ | Circadian driving frequency |
| $T_{\text{sim}}$ | 365 d | Stochastic simulation duration (1 year) |
| $\alpha$ | 0-1 for scan;<br>0.5 or 0.3 in examples | Circadian amplitude |
| $\theta$ | 0.3 | OU mean-reversion rate in model-time units |
| $\sigma_\eta$ | 0.006, 0.010, 0.012, 0.015 | OU perturbation intensities in occupancy scans |

### 3.2. Circadian forcing confines trajectories to the physiological pair

Circadian input acts on the system according to the slow-fast logic (**Fig. 1B**). It shifts the fast nullcline, creating and destroying fixed points on the slow nullcline and thereby biasing attractor choice. In the potential representation, this shift appears as a tilt of the fast potential (**Fig. 1C**). In the fixed amplitude regime, this tilt is sufficient to open the inner physiological transition between L and H while avoiding sustained entry into PL or PH over the short observation window. Consequently, the trajectory alternates daily between physiological L and H without perturbations, and low-intensity OU perturbations add fluctuations around the same circadian path without sustained outer-well dwell time (**Fig. 2**).

Circadian input can switch the affective state by tilting the effective potential (**Fig. 3**). In a generic landscape (*a*_3_ = 1.593), the fast potential is evaluated at three slow-coordinate slices near the physiological L state, the middle saddle, and the physiological H state. At the L slice, the high-input tilt lowers the potential toward larger *y*, favoring motion toward H. At the middle saddle, small changes in tilt can select either side of the physiological pair. At the H slice, the low-input tilt lowers the potential toward smaller *y*, favoring motion back toward L. With OU perturbations, the affective state can escape past the barriers into a lower potential well. Circadian forcing alternates the favored direction of motion within the inner physiological pair, rather than acting as a biased stochastic drive.

### 3.3. Analytical stability criterion

Eq. (19) converts the phase-dependent rate approximation into a per-cycle escape hazard, and Eq. (20) converts that hazard into a finite-horizon escape probability. The per-cycle hazard *H_p_*_→*d*_ measures the expected pathological escape count during one circadian cycle, so *H_p_*_→*d*_ ≪ 1 indicates rare escape per cycle. However, the long stochastic simulations span many cycles; Eq. (20) shows that sustained confinement over the full observation window requires *NH_p_*_→*d*_ ≪ 1, not merely *H_p_*_→*d*_ ≪ 1. Mechanistically, changing circadian amplitude reshapes the phase-dependent barrier hierarchy and the time spent near each basin. In the simulated geometries, reduced amplitude weakens regular physiological L-H entrainment and increases occupancy near pathological-access regions, while increasing OU intensity *σ_η_* or decreasing mean-reversion rate *θ* increases the effective perturbation scale *D*_eff_ in Eq. (18). The present simulations vary perturbation intensity *σ_η_* at fixed *θ*; the predicted effect of longer OU persistence remains to be tested in a dedicated *θ* scan. The final dwell-time statistics are computed directly from one-year stochastic trajectories rather than from an additional two-state hazard formula.

### 3.4. One-year occupancy regimes

We traced representative one-year trajectories for three landscape geometries that phenomenologically resemble distinct affective regimes and produce different occupancy patterns as the central nullcline bend *a*_3_ changes (**Figs. 4**-**6**). In the mania-prone contrast (*a*_3_ = 1.595), PH occupancy is appreciable under the same zero-mean OU perturbation protocol, especially when circadian amplitude is reduced (**Fig. 4**). In the euthymia-like geometry (*a*_3_ = 1.605), occupancy is mostly physiological across the preserved-amplitude examples, with reduced circadian amplitude increasing PL occupancy but not producing a permanently absorbing state (**Fig. 5**). In the BD-I-like proxy geometry (*a*_3_ = 1.620), PH occupancy remains reachable but rare, whereas PL occupancy is more common and physiological state occupancy remains substantial (**Fig. 6**). This pattern is consistent with finite-horizon pathological entry followed by nonabsorbing but prolonged outer-well dwell time.

**Figure 5.**
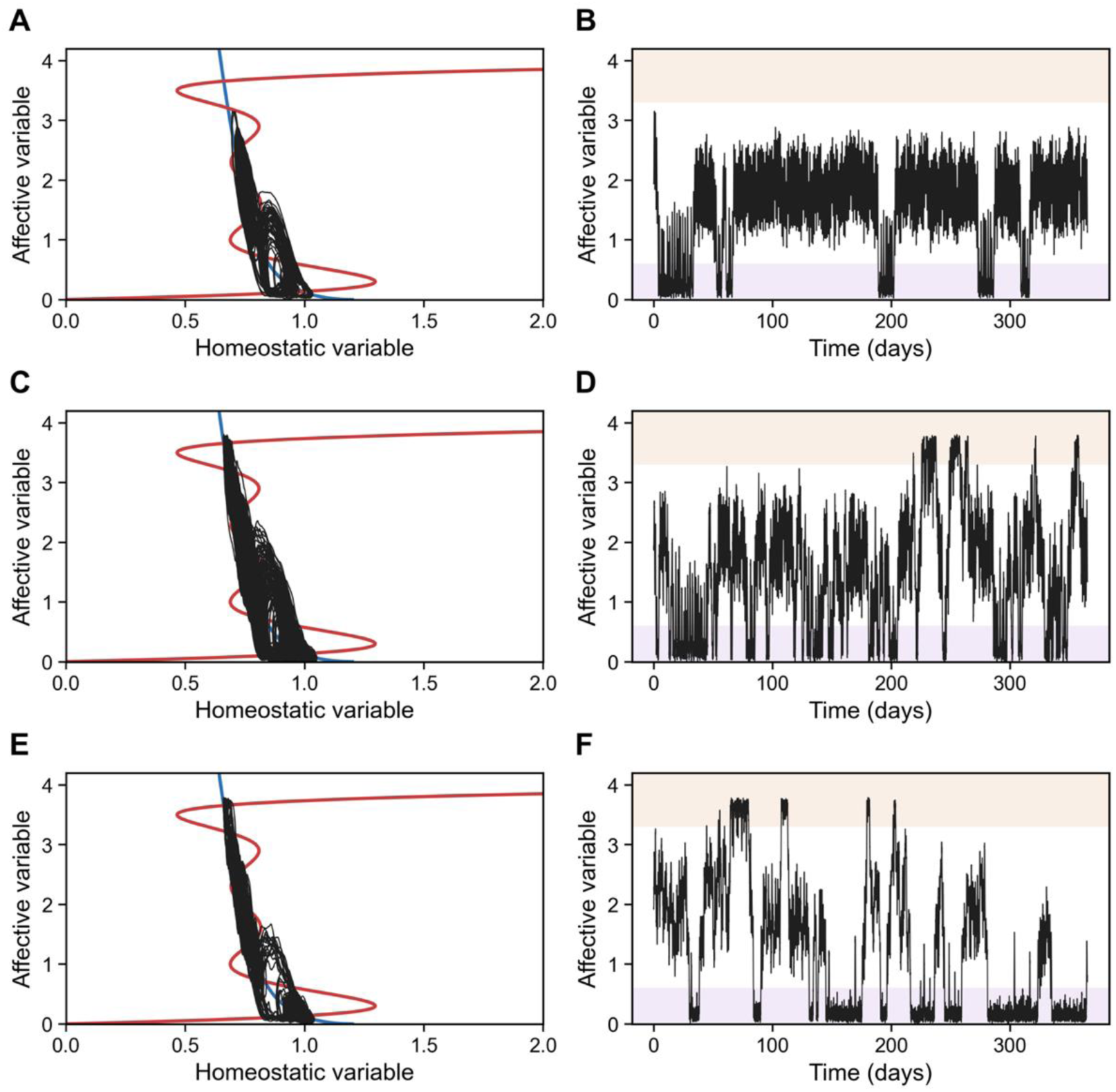
Representative one-year transitions in the euthymia-like landscape. Simulations use the euthymia-like geometry (*a*_3_ = 1.605, *θ* = 0.3). The left column (**A,C,E**) shows phase-plane trajectories over the deterministic nullclines, and the right column (**B,D,F**) shows the corresponding fast affective coordinate over 365 days. (**A,B**) Preserved circadian amplitude (*α* = 0.5) with low OU perturbation intensity (*σ_η_* = 0.006) produces normal alternations predominantly within the inner physiological wells (L and H), with only brief PL dips. (**C,D**) Preserved amplitude (*α* = 0.5) with stronger perturbation (*σ_η_* = 0.012) produces larger recurrent excursions with transient access to the pathological regions, although the trajectory repeatedly returns to the physiological range. (**E,F**) Reduced amplitude (*α* = 0.3) with the same stronger perturbation increases occupancy in the PL state, but the trajectory remains recurrent rather than permanently absorbing.

**Figure 6.**
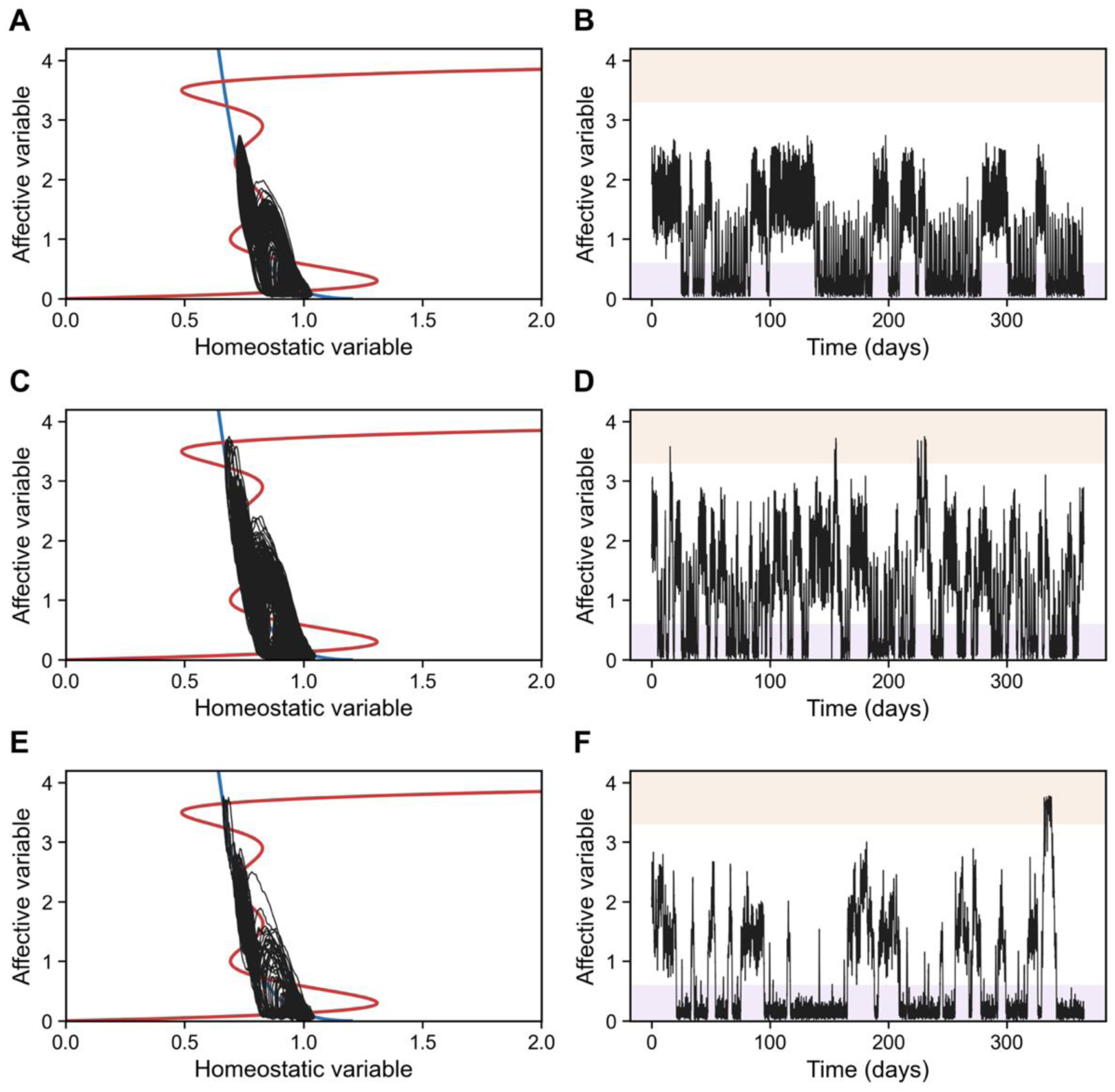
Representative one-year transitions in the BD-I-like proxy landscape. Simulations use the BD-I-like geometry (*a*_3_ = 1.620, *θ* = 0.3). The left column (**A,C,E**) shows phase-plane trajectories over the deterministic nullclines, and the right column (**B,D,F**) shows the corresponding fast affective coordinate over 365 days. (**A,B**) Preserved circadian amplitude (*α* = 0.5) with low OU perturbation intensity (*σ_η_* = 0.006) produces mostly physiological state occupancy with substantial PL visits and no PH occupancy in this representative trajectory. (**C,D**) Preserved amplitude (*α* = 0.5) with stronger perturbation (*σ_η_* = 0.012) maintains substantial physiological state occupancy and admits rare PH excursions. (**E,F**) Reduced amplitude (*α* = 0.3) with the same stronger perturbation increases total pathological dwell time, dominated by the PL region, while retaining limited access to the PH region.

Small changes in the central nullcline bend *a*_3_ therefore bias occupancy statistics without changing the OU perturbation mean. Decreasing *a*_3_ to 1.595 produces a mania-prone polarity-bias contrast, whereas the intermediate geometry *a*_3_ = 1.605 is relatively euthymia-like in the sense of mostly combined physiological state occupancy. The *a*_3_ = 1.620 case serves as a BD-I-like proxy geometry because it combines substantial physiological state occupancy with rare but nonzero PH occupancy and greater PL burden. These simulations should not be interpreted as fitting a specific temperament or diagnostic subtype. They show that a small geometric asymmetry can convert the same four-well mechanism into different occupancy regimes.

### 3.5. Parameter scans and fractional occupancy

**Fig. 7** summarizes the parameter scans for aggregate occupancy statistics corresponding to the representative examples. We first confirm that the three representative values of the central nullcline bend *a*_3_ lie inside the four-well region of the parameter slice (**Fig. 7A**). We then hold the OU parameters fixed (*σ_η_* = 0.012, *θ* = 0.3) and probe how changing circadian amplitude redistributes occupancy among PL, PH, and the combined physiological pair (L+H) to emphasize outer-well burden (**Fig. 7B**). These traces are averaged across eight independent stochastic realizations per parameter value. For the BD-I-like geometry, the stochastic occupancy percentages over one year are computed for PL, L, H, and PH regimes as *α* and *σ_η_* vary, averaged across 30 independent realizations per parameter value (**Fig. 7C**). The mania-prone geometry retains appreciable PH occupancy over the scan, the euthymia-like geometry has the largest mostly physiological range, and the BD-I-like geometry shows rare but nonzero PH occupancy with a larger PL burden. It also shows how one-year fractional occupancy redistributes across physiological and pathological regions as perturbation intensity increases. At *α* = 0.5 and *σ_η_* = 0.012, the trajectory ensemble spends approximately 36% of the year in PL, 1% in PH, and 63% in the physiological pair. Reducing *α* to 0.3 at the same OU intensity increases PL occupancy to about 59%, while PH remains below 1%. In this BD-I-like proxy, PH occupancy is reachable but rare and PL occupancy accounts for most episode-like burden in the BD-I-like candidate over the one-year window.

**Figure 7.**
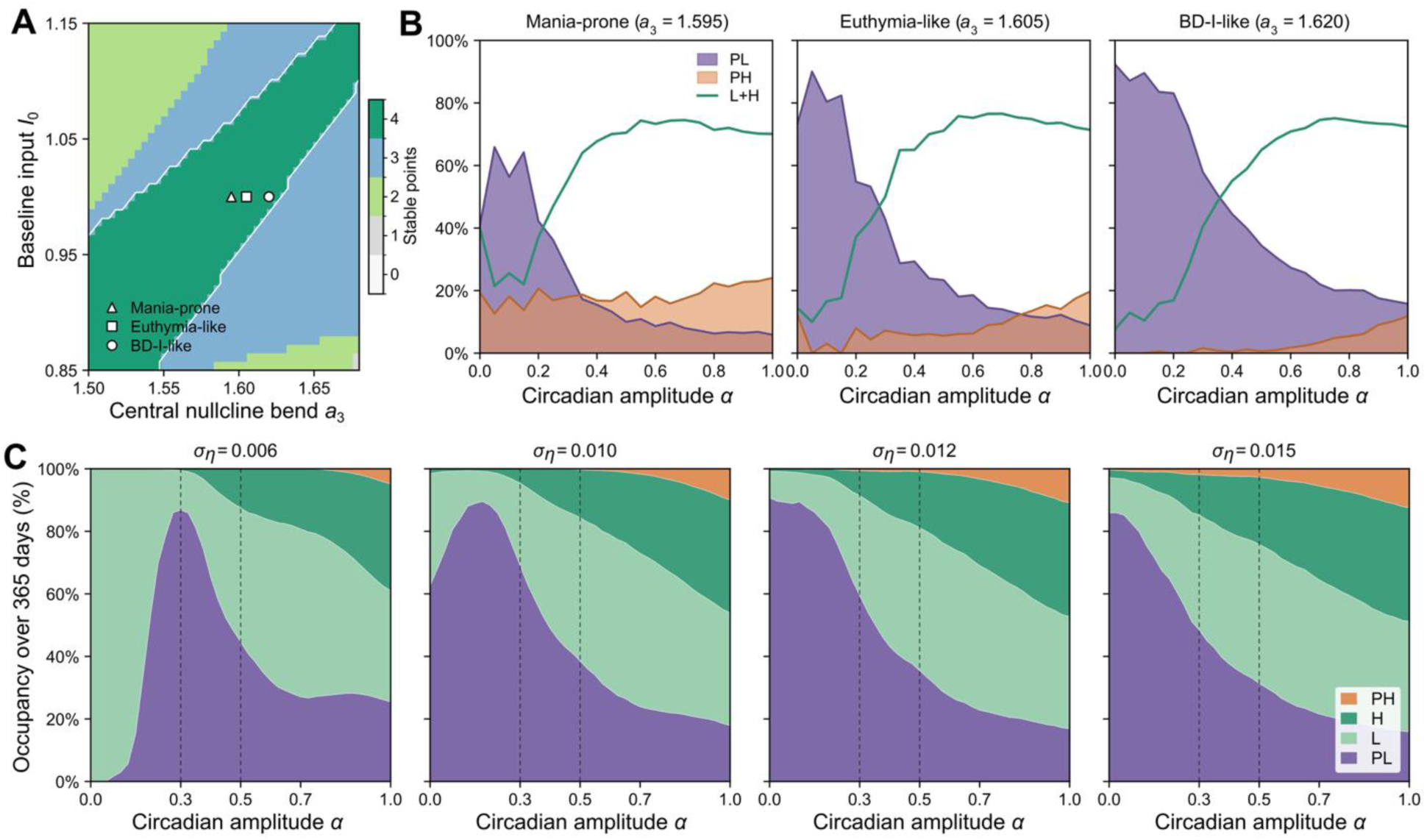
Topology, geometry-dependent occupancy regimes, and OU perturbation-intensity dependence. (**A**) Topology map of the number of stable fixed points over the (*a*_3_, *I*_0_) plane; the color bar gives the count. Markers (triangle, square, circle) indicate the three representative geometries used in the occupancy analyses, all at baseline input *I*_0_ = 1.0: mania-prone (*a*_3_ = 1.595), euthymia-like (*a*_3_ = 1.605), and BD-I-like (*a*_3_ = 1.620). (**B**) Geometry-dependent circadian-amplitude scans at fixed OU perturbation intensity *σ_η_* = 0.012, mean-reversion rate *θ* = 0.3, and baseline input *I*_0_ = 1.0. Traces give the percentage of a 365-day simulation spent in the PL and PH regions and in the combined physiological pair (L+H). The euthymia-like geometry holds the highest physiological occupancy at large amplitude, whereas the mania-prone and BD-I-like geometries carry different pathological-polarity burdens under the same zero-mean OU protocol. (**C**) Stacked occupancy percentages for the BD-I-like geometry (*a*_3_ = 1.620) as OU perturbation intensity varies across *σ_η_* = 0.006, 0.010, 0.012, and 0.015. Colors give the PH, H, L, and PL fractions. Dashed vertical lines mark the representative circadian amplitudes *α* = 0.3 and *α* = 0.5 used in the trajectory examples of Figs. 4-6. The labels mania-prone, euthymia-like, and BD-I-like denote model occupancy patterns, not clinical diagnoses.

### 3.6. Circadian phase reshapes potential accessibility

The full circadian phase-dependent accessibility of the broader landscape is summarized in **Fig. 8**. The circadian mechanism underlying **Figs. 2** and **7** is shown with the same circadian input at *α* = 0.3 and *α*=0.5 as a phase-dependent potential tilt at fixed slow coordinate *x*. In the time-series view, *I*(*t*) is an oscillatory input. In the fixed-*x* fast-potential view, the same input appears as the phase-dependent linear term −*γI*(*t*)*x*_0_*y* in Eq. (15). Changing circadian phase *φ* therefore changes the relative barrier heights of the wells rather than applying a separate force outside the landscape. This fixed-*x* view is not a full trajectory calculation; rather, it visualizes the instantaneous accessibility structure that contributes to the fractional occupancy measured in the stochastic simulations.

**Figure 8.**
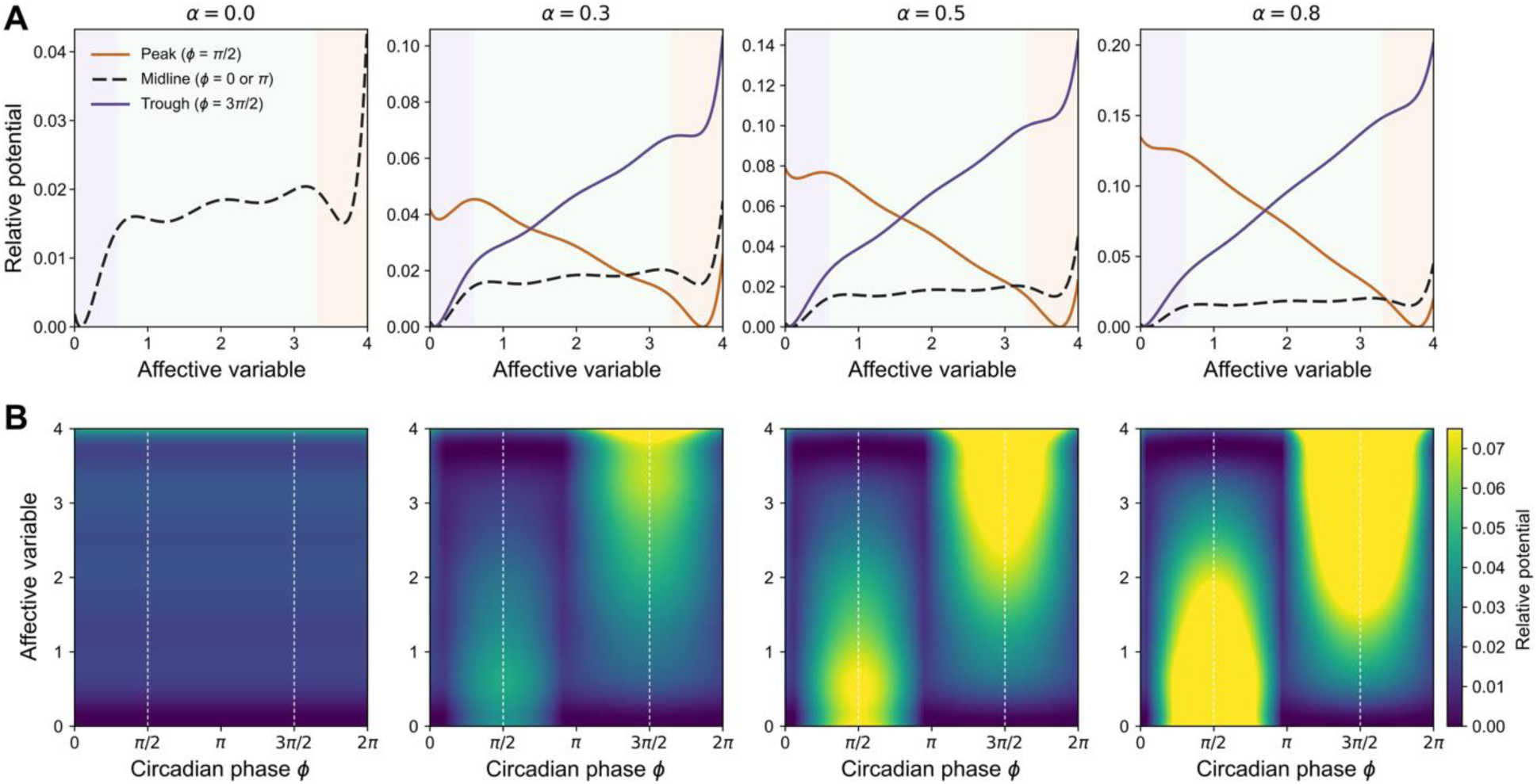
Circadian input as a phase-dependent tilt of the fast affective potential. The circadian oscillatory input *I*(*φ*) enters the fast potential as the linear tilt term −*γ I*(*φ*)*x*_0_*y*, evaluated here at fixed slow coordinate *x* = *x*_0_ = 0.738. All curves and heatmaps show the relative potential, and the background shading follows the PL, physiological, and PH convention of Fig. 1. (**A**) Representative potential profiles at circadian peak (*φ* = *π*/2), midline (*φ* = 0 or *π*), and trough (*φ* = 3*π*/2) for circadian amplitudes *α* = 0, 0.3, 0.5, and 0.8. At *α* = 0, the phase-dependent tilt vanishes, so the phase profiles coincide and only the midline profile is shown. For nonzero *α*, the peak and trough phases tilt the potential in opposite directions and change the relative depths of the low- and high-affective regions. (**B**) Heatmaps of the relative potential over circadian phase and affective coordinate for the same *α* values. Vertical dashed lines mark peak input (*φ* = *π*/2) and trough input (*φ* = 3*π*/2). Increasing *α* strengthens the phase-dependent relocation of low-potential regions: circadian forcing can open physiological transition windows, while larger tilts can also increase access to the outer pathological wells.

**Fig. 8A** shows representative potential profiles at circadian peak, midline, and trough. When *α* = 0, the three phases coincide because the tilt is absent. When *α* > 0, peak and trough phases tilt the potential in opposite directions; because the plotted quantity is the relative potential, this appears as alternating elevation of one side of the finite *y* interval. **Fig. 8B** extends this view across the full circadian phase. As *α* increases, the low-potential valley shifts more strongly with phase and local wells can become transiently less accessible in the fixed-*x* cross-section. This is the graphical counterpart of the analytical hazard: circadian amplitude controls which barriers are opened for physiological switching, while the outer pathological barriers remain protected only over a finite range of geometry and perturbation intensity.

## 4. DISCUSSION

Here, we present a biologically motivated yet phenomenological model that illustrates a possible dynamical mechanism by which weakened circadian rhythms and persistent stochastic load increase vulnerability to prolonged pathological states, measured as dwell time or fractional occupancy across both clinical episodes and normal mood states. Although the model is motivated by the slow-fast separation of HPA-axis homeostatic feedback, we extend the potential landscape to multiple wells and consider the role of circadian tilt, which helps keep the duration and intensity of affective states within a normal homeostatic range of fluctuation. This model should be understood as a biologically motivated but phenomenological slow-fast affective landscape that provides a dynamical mechanism rather than as a direct mapping from hormone concentrations to mood states. This framing is consistent with the dynamical-disease perspective, in which disease-relevant behavior can emerge from altered feedback dynamics and changes in qualitative system organization rather than from a single static abnormality (Mackey and Glass, 1977; Glass and Mackey, 1979; Mackey and Milton, 1987).

In our model, circadian rhythm alterations interact with sensitivity to environmental perturbations (broadly defined) to drive transitions into pathological mood states. The sensitivity parameter represents an intrinsic property of the system, or an individual-level susceptibility, rather than the external inputs themselves. This framing is consistent with the clinical understanding that bipolar disorder is fundamentally a brain disorder and that external stressors act as triggers that precipitate psychopathological states in an already altered neural system (Lex et al., 2017). More specifically, the model suggests that heightened sensitivity to environmental changes triggers transient pathological states, whereas intrinsic circadian disruption provides a baseline vulnerability that stabilizes these states. Across the affective spectrum, this division suggests a possible interpretation of distinct clinical phenotypes. A predominance of hypersensitivity might manifest in conditions characterized by mood instability and rapid, transient mood shifts, such as cyclothymia or borderline personality disorder (American Psychiatric Association, 2022). In contrast, a predominance of intrinsic circadian disruption might underlie conditions characterized by more sustained manic or depressive episodes that are relatively insensitive to external factors, as typically observed in BD-I (Perugi et al., 2011; Perugi et al., 2015).

Acute and chronic stress are commonly implicated in mood disorders; for example, patients with bipolar disorder or major depressive disorder often experience mood episodes following environmental or social stressors (Lex et al., 2017). These stressors are represented as OU perturbations in our model. The OU process is not introduced because white noise cannot produce transitions. White noise could produce qualitatively similar rare escapes in some regimes. Instead, OU noise is used because stressor load represents perturbations with finite autocorrelation rather than independent instantaneous impulses. The present simulations vary perturbation intensity *σ_η_* at fixed *θ* = 0.3; Eq. (18) predicts that longer OU persistence, corresponding to smaller *θ*, would similarly increase the effective perturbation scale.

Our model suggests that high sensitivity to stressors, rather than the stressors themselves, plays an important role in pathological mood shifts. The landscape geometry provides a conceptual framework for considering how circadian modulation and perturbation sensitivity jointly govern episode vulnerability. With preserved circadian amplitude, correlated fluctuations can produce transient excursions that typically return to the physiological pair (**Fig. 2**). In contrast, reduced circadian amplitude weakens regular low-high entrainment and, when combined with stronger or more persistent stochastic load, increases noise-driven escape probability and prolongs dwell times in outer pathological wells. This provides a dynamical explanation for how the same circadian mechanism that normally organizes daily affective variation may, when weakened, lose its protective role against episode-scale dwell time in depression-like or mania-like states.

We also examined transitions to pathological mood states under geometrically biased landscapes. Simulations show that dominant affective polarity is controlled primarily by the central nullcline-bend parameter *a*_3_, while circadian amplitude *α* and OU perturbation intensity *σ_η_* regulate escape and occupancy over the one-year window (**Figs. 4**-**6**). Slight variation in *a*_3_ preferentially changes access to PH or PL rather than symmetrically increasing access to both. Varying circadian amplitude and sensitivity to external noise yields patterns similar to those of the unbiased system, although transitions are primarily observed in one direction. By tuning *a*_3_, the model spans a continuum ranging from a strong preference for the PH state (mania-prone) to a strong preference for the PL state, including intermediate regimes such as the euthymia-like geometry and the BD-I-like proxy geometry, with rare but reachable PH state occupancy and a larger PL burden (**Fig. 7**).

This mechanism offers a possible dynamical explanation for why depressive states can predominate longitudinally even in BD-I. Bipolar disorder includes clinically distinct but related course patterns. BD-I is defined by the occurrence of mania, whereas BD-II involves hypomanic and depressive episodes without full mania, and cyclothymic disorder involves chronic subthreshold hypomanic and depressive symptoms. In the model, this asymmetry does not require biased stochastic forcing; instead, it can emerge from the geometric alignment of the slow homeostatic nullcline with the fast affective nullcline. A relatively steep or vertically aligned slow nullcline may be interpreted as a form of homeostatic inertia, in which the slow regulatory coordinate changes less readily than the fast affective coordinate. In the original HPA-axis framework, the slow component corresponds to reserve CRH regulated by feedback, whereas acute perturbations act immediately through circulating CRH release. By analogy, in our affective landscape formulation, the homeostatic coordinate can be viewed as a slow regulatory constraint that resists rapid reconfiguration after perturbation, naturally producing depression-biased dwell time while still allowing rare access to mania-like states. This may reflect physiological conservatism in response to external perturbations. Clinically, among the multiple factors thought to underlie the dominance of manic or depressive polarity, temperament has long been considered to provide biological predisposition as part of the soft bipolar spectrum (Favaretto et al., 2024). Affective-temperament models are themselves asymmetric in that hyperthymic temperament represents the clearest high-energy, mania-adjacent disposition, whereas depressive, anxious, cyclothymic, and irritable temperaments are more closely aligned with negative affect, withdrawal, affective instability, or dysregulated reactivity. This temperamental asymmetry is paralleled by longitudinal studies of bipolar disorder showing that depressive symptoms occupy substantially more time than manic or hypomanic symptoms across the illness course (Judd et al., 2002).

The modeling results generate testable hypotheses about circadian rhythm disturbances in bipolar disorder. Circadian rhythm alternations, including decreased amplitude, increased variability, and phase advances or delays, are commonly observed in bipolar disorder (Esaki et al., 2021), although they are generally considered peripheral to the core symptomatology of manic-depressive psychopathology (American Psychiatric Association, 2022). Seasonality is well documented in patients with depressive and bipolar disorders. Typically, depressive episodes tend to begin in autumn or winter and remit in spring, whereas manic or hypomanic episodes more often peak in spring and summer (Geoffroy et al., 2014). Seasonal variation in photoperiod and light exposure also reshapes circadian and sleep parameters, with reduced winter light linked to weaker and less regular circadian organization (Skeldon et al., 2022). Within our framework, seasonality enters as slow annual modulation of the circadian drive. The circadian amplitude *α* can drift with photoperiod across the year (Schmal et al., 2020). A seasonal reduction in *α* weakens periodic confinement to the physiological pair and raises dwell time in the outer wells, lengthening pathological dwell time without by itself favoring one polarity. A seasonal shift in the baseline *I*_0_ sets the predominant polarity: a short winter photoperiod that lowers mean light input biases the tilt toward the low wells, whereas a long summer photoperiod biases it toward the high wells, while the depth and asymmetry of the wells remain governed by the central bend parameter *a*_3_ (**Figs. 7** and **8**). The reverse seasonal pattern of winter depression and summer mania therefore maps onto a slow annual oscillation in baseline drive superimposed on the daily tilt that gates physiological switching. Seasonality is, in this sense, a natural slow-modulation extension of the present model, in which the same circadian input carries fast daily and slow annual components that together shift fractional occupancy across the four wells.

The biological substrates of circadian rhythm disruption and hypersensitivity to environmental factors remain to be clarified. At the behavioral level, our model may be used to describe psychomotor activity, which is closely linked to mood and energy. Psychomotor activity can be objectively measured via actigraphy and is considered a core dimension of psychopathology in bipolar disorder: it is increased during manic states, manifesting as hyperactivity and impulsivity and often accompanied by euphoria or dysphoria, and decreased in typical depressive states, where it presents as psychomotor retardation and is frequently associated with anhedonia (Faurholt-Jepsen et al., 2016; Martino & Magioncalda, 2024). At the biological level, the model could also be extended to represent neurotransmitter dynamics. Dopamine, a neurotransmitter critically involved in motor control and motivation, as well as serotonin, have been implicated in a wide range of psychiatric disorders, including bipolar disorder. Dopaminergic signaling has been proposed to increase during manic states and decrease during depressive states, whereas serotonergic signaling is generally thought to be reduced during mania (Conio et al., 2020; Magioncalda & Martino, 2022; Martino & Magioncalda, 2022; Cabana-Domínguez et al., 2022). Notably, dopamine expression exhibits strong circadian oscillations across multiple brain regions and is regulated by core clock genes, and dopamine can also feed back onto clock gene regulation (Korshunov et al., 2017; Rahim et al., 2025). Dopamine-dependent arousal dynamics may be mediated by an intrinsic ultradian oscillator that normally interacts with the circadian clock but can desynchronize and undergo marked period lengthening under elevated dopamine tone, reaching infradian timescales in sleep-wake and heightened-activity behaviors relevant to bipolar cycling (Blum et al., 2014; Markam et al., 2025). Finally, our model could be extended to describe brain network dynamics. Intrinsic brain activity and network balance exhibit opposing alterations across manic and depressive states: during mania, intrinsic activity within the sensorimotor network is increased, or the balance between the sensorimotor and default-mode networks shifts toward sensorimotor dominance, potentially reflecting immediate interaction with the external environment; conversely, during depressive states, intrinsic activity within the sensorimotor network is reduced, or the network balance shifts toward default-mode dominance, potentially reflecting detachment from the external environment (Martino et al., 2016; Magioncalda & Martino, 2022; Mavar et al., 2025). Our landscape model provides a natural platform for integrating these phenomena by allowing the tilt or geometry of the landscape, for example through the parameter *a*_3_, to be influenced by additional oscillatory components operating on different timescales.

Several limitations clarify scope and motivate extensions. The seventh-order nullcline is a phenomenological choice that enforces quadristability; although the minimal-polynomial argument justifies why a septic form is mathematically sufficient for four wells, it does not identify the biological source of the four-well topology. The current simulations vary circadian disruption primarily through amplitude; phase shifts, day-to-day variability, light exposure history, and altered waveform shape may yield distinct transition statistics. Finally, the model could be extended to include additional biological processes operating on distinct timescales. Dopamine-based arousal oscillations or resting-state network reconfigurations could introduce perturbations on the slow coordinate; circadian and seasonal inputs could be coupled more directly to endocrine dynamics; and slower forms of endocrine adaptation operating on multiday timescales could be incorporated (Karin et al., 2020). Despite these limitations, the framework provides testable predictions. It predicts that reduced circadian amplitude and increased perturbation sensitivity will increase the frequency of escapes into extreme states and the dwell-time asymmetry between manic-like and depressive-like attractors under polarity bias. It also predicts that interventions that strengthen circadian amplitude or reduce effective noise correlation should restore robust alternation within the physiological pair and reduce pathological dwell time.

## 5. CONCLUSION

We developed a quadristable slow-fast model of affective-state switching under circadian drive. The model treats circadian input as a global oscillatory tilt of a multistable affective landscape and shows how daily alternations between physiological low and high states, as well as irregular, infradian pathological episodes, can arise from the interaction among circadian amplitude, persistent perturbation load, and landscape geometry. Fractional occupancy analyses provide an interpretable bridge between dynamical regimes and bipolar-spectrum course patterns while preserving the distinction between model-proxy phenotypes and clinical diagnosis.

## ACKNOWLEDGMENTS

V.W. received support from the Fulbright U.S. Student Program, sponsored by the U.S. Department of State and the Foundation for Scholarly Exchange (Fulbright Taiwan). P.M., M.M., and J.M. received support from the National Science and Technology Council (NSTC), Taiwan (P.M.: 113-2314-B-038-096-MY3; M.M.: 113-2628-B-038-010-MY3; J.M.: 107-2410-H-038-004-MY2, 113-2314-B-038-121, 114-2320-B-038-052-MY3). J.M. also received support from the Higher Education Sprout Project of the Ministry of Education (MOE), Taiwan.

## AUTHOR DECLARATIONS

### Author Contributions

V.W.: Conceptualization, formal analysis, investigation, visualization, validation, methodology, writing - original draft, writing - review and editing. P.M.: Conceptualization, clinical interpretation, writing - review and editing. M.M.: Conceptualization, clinical interpretation, writing - review and editing. J.M.: Conceptualization, formal analysis, investigation, visualization, validation, methodology, supervision, writing - original draft, writing - review and editing.

## DATA AVAILABILITY

The simulation code and generated trajectory data are available from the corresponding author upon reasonable request.

## REFERENCES

Adam EK, Quinn ME, Tavernier R, McQuillan MT, Dahlke KA, Gilbert KE (2017). Diurnal cortisol slopes and mental and physical health outcomes: A systematic review and meta-analysis. Psychoneuroendocrinology, 83:25–41.

American Psychiatric Association (2022). Diagnostic and statistical manual of mental disorders (5th ed., text rev.). 10.1176/appi.books.9780890425787

Andersen M, Vinther F, Ottesen JT (2013). Mathematical modeling of the hypothalamic-pituitary-adrenal gland (HPA) axis, including hippocampal mechanisms. Math Biosci, 246:122–138.

Blum ID, Zhu L, Moquin L, Kokoeva MV, Gratton A, Giros B, Storch KF (2014). A highly tunable dopaminergic oscillator generates ultradian rhythms of behavioral arousal. eLife, 3:e05105.

Cabana-Domínguez J, Torrico B, Reif A, Fernàndez-Castillo N, Cormand B (2022). Comprehensive exploration of the genetic contribution of the dopaminergic and serotonergic pathways to psychiatric disorders. Transl Psychiatry, 12:11.

Carvalho AF, Firth J, Vieta E (2020). Bipolar disorder. N Engl J Med, 383:58–66.

Cheng X, D’Orsogna MR, Chou T (2021). Mathematical modeling of depressive disorders: Circadian driving, bistability and dynamical transitions. Comput Struct Biotechnol J, 19:664–690.

Conio B, Martino M, Magioncalda P, Escelsior A, Inglese M, Amore M, Northoff G (2020). Opposite effects of dopamine and serotonin on resting-state networks: review and implications for psychiatric disorders. Mol Psychiatry, 25:82–93.

Daugherty D, Roque-Urrea T, Urrea-Roque J, Troyer J, Wirkus S, Porter MA (2009). Mathematical models of bipolar disorder. Commun Nonlinear Sci Numer Simul, 14:2897–2908.

Dollish HK, Tsyglakova M, McClung CA (2024). Circadian rhythms and mood disorders: Time to see the light. Neuron, 112:25–40.

Esaki Y, Obayashi K, Saeki K, Fujita K, Iwata N, Kitajima T (2021). Association between circadian activity rhythms and mood episode relapse in bipolar disorder: a 12-month prospective cohort study. Transl Psychiatry, 11:525.

Faurholt-Jepsen M, Brage S, Vinberg M, Kessing LV (2016). State-related differences in the level of psychomotor activity in patients with bipolar disorder–Continuous heart rate and movement monitoring. Psychiatry Res, 237:166–174.

Favaretto E, Bedani F, Brancati GE, De Berardis D, Giovannini S, Scarcella L, Martiadis V, Martini A, Pampaloni I, Perugi G, Pessina E (2024). Synthesising 30 years of clinical experience and scientific insight on affective temperaments in psychiatric disorders: State of the art. J Affect Disord, 362:406–415.

Geoffroy PA, Bellivier F, Scott J, Etain B (2014). Seasonality and bipolar disorder: a systematic review, from admission rates to seasonality of symptoms. J Affect Disord, 168:210–223.

Glass L, Mackey MC (1979). Pathological conditions resulting from instabilities in physiological control systems. Ann N Y Acad Sci, 316:214–235.

Goldbeter A (2011). A model for the dynamics of bipolar disorders. Prog Biophys Mol Biol, 105:119–127.

Grande I, Berk M, Birmaher B, Vieta E (2016). Bipolar disorder. Lancet, 387:1561–1572.

Gupta S, Aslakson E, Gurbaxani BM, Vernon SD (2007). Inclusion of the glucocorticoid receptor in a hypothalamic pituitary adrenal axis model reveals bistability. Theor Biol Med Model, 4:8.

Herane-Vives A, Arnone D, De Angel V, Papadopoulos A, Wise T, Alameda L, Chua KC, Young AH, Cleare AJ (2020). Cortisol levels in unmedicated patients with unipolar and bipolar major depression using hair and saliva specimens. Int J Bipolar Disord, 8:15.

Huang MC, Chuang SC, Tseng MC, Chien YL, Liao SC, Chen HC, Kuo PH (2017). Cortisol awakening response in patients with bipolar disorder during acute episodes and partial remission: A pilot study. Psychiatry Res, 258:594–597.

Inder ML, Crowe MT, Porter R (2016). Effect of transmeridian travel and jetlag on mood disorders: Evidence and implications. Aust N Z J Psychiatry, 50:220–227.

Jelić S, Čupić Ž, Kolar-Anić L (2005). Mathematical modeling of the hypothalamic-pituitary-adrenal system activity. Math Biosci, 197:173–87.

Jones JR, Chaturvedi S, Granados-Fuentes D, Herzog ED (2021). Circadian neurons in the paraventricular nucleus entrain and sustain daily rhythms in glucocorticoids. Nat Commun, 12:5763.

Judd LL, Akiskal HS, Schettler PJ, Endicott J, Maser J, Solomon DA, Leon AC, Rice JA, Keller MB (2002). The long-term natural history of the weekly symptomatic status of bipolar I disorder. Arch Gen Psychiatry, 59:530–537.

Karin O, Raz M, Tendler A, Bar A, Korem Kohanim Y, Milo T, Alon U (2020). A new model for the HPA axis explains dysregulation of stress hormones on the timescale of weeks. Mol Syst Biol, 16:e9510.

Kim LU, D’Orsogna MR, Chou T (2016). Onset, timing, and exposure therapy of stress disorders: mechanistic insight from a mathematical model of oscillating neuroendocrine dynamics. BMC Biol Direct, 11:13.

Korshunov KS, Blakemore LJ, Trombley PQ (2017). Dopamine: A Modulator of Circadian Rhythms in the Central Nervous System. Front Cell Neurosci, 11:91.

Lee A, Myung SK, Cho JJ, Jung YJ, Yoon JL, Kim MY (2017). Night shift work and risk of depression: Meta-analysis of observational studies. J Korean Med Sci, 32:1091–1096.

Lex C, Baezner E, Meyer TD (2017). Does stress play a significant role in bipolar disorder? A meta-analysis. J Affect Disord, 208:298–308.

Mackey MC, Glass L (1977). Oscillation and chaos in physiological control systems. Science, 197:287–289.

Mackey MC, Milton JG (1987). Dynamical diseases. Ann N Y Acad Sci, 504:16–32.

Magioncalda P, Martino M (2022). A unified model of the pathophysiology of bipolar disorder. Mol Psychiatry, 27:202–211.

Maripuu M, Wikgren M, Karling P, Adolfsson R, Norrback KF (2014). Relative hypo-and hypercortisolism are both associated with depression and lower quality of life in bipolar disorder: a cross-sectional study. PLOS One, 9:e98682.

Markam PS, Bourguignon C, Zhu L, Ward B, Darvas M, Sabatini PV, Kokoeva MV, Giros B, Storch KF (2025). Mesolimbic dopamine neurons drive infradian rhythms in sleep-wake and heightened activity state. Sci Adv, 11:eado9965.

Martino M, Magioncalda P, Huang Z, Conio B, Piaggio N, Duncan NW, Rocchi G, Escelsior A, Marozzi V, Wolff A, Inglese M (2016). Contrasting variability patterns in the default mode and sensorimotor networks balance in bipolar depression and mania. Proc Natl Acad Sci USA, 113:4824–9.

Martino M, Magioncalda P (2022). Tracing the psychopathology of bipolar disorder to the functional architecture of intrinsic brain activity and its neurotransmitter modulation: a three-dimensional model. Mol Psychiatry, 27:793–802.

Martino M, Magioncalda P (2024). A three-dimensional model of neural activity and phenomenal-behavioral patterns. Mol Psychiatry, 29:639–652.

Mavar S, Lee YS, Baranova E, Duncan NW, Magioncalda P, Martino M (2025). A working model linking the psychopathology and pathophysiology of major depressive disorder-an umbrella review of neuroimaging studies and a conceptual framework. Mol Psychiatry, 30:6007–6019.

McIntyre RS, Berk M, Brietzke E, Goldstein BI, López-Jaramillo C, Kessing LV, Malhi GS, Nierenberg AA, Rosenblat JD, Majeed A, Vieta E (2020). Bipolar disorders. Lancet, 396:1841–1856.

Milo T, Maimon L, Cohen B, Haran D, Segman D, Danon T, Bren A, Mayo A, Rappaport GC, McInnis M, Alon U (2024). Longitudinal hair cortisol in bipolar disorder and a mechanism based on HPA dynamics. iScience, 27:109234.

Murri MB, Prestia D, Mondelli V, Pariante C, Patti S, Olivieri B, Arzani C, Masotti M, Respino M, Antonioli M, Vassallo L (2016). The HPA axis in bipolar disorder: systematic review and meta-analysis. Psychoneuroendocrinology, 63:327–42.

Nunes A, Singh S, Allman J, Becker S, Ortiz A, Trappenberg T, Alda M (2022). A critical evaluation of dynamical systems models of bipolar disorder. Transl Psychiatry, 12:146.

Perugi G, Fornaro M, Akiskal HS (2011). Are atypical depression, borderline personality disorder and bipolar II disorder overlapping manifestations of a common cyclothymic diathesis? World Psychiatry, 10:45.

Perugi G, Hantouche E, Vannucchi G, Pinto O (2015). Cyclothymia reloaded: A reappraisal of the most misconceived affective disorder. J Affect Disord, 183:119–33.

Rahim AR, Will V, Myung J (2025). Mood variation under dual regulation of circadian clock and light. Chronobiol Int, 42:162–184.

Schmal C, Herzel H, Myung J (2020). Clocks in the wild: entrainment to natural light. Front Physiol, 11:272.

Skeldon AC, Dijk DJ, Meyer N, Wulff K (2022). Extracting circadian and sleep parameters from longitudinal data in schizophrenia for the design of pragmatic light interventions. Schizophr Bull, 48:447–456.

Steinacher A, Wright KA (2013). Relating the bipolar spectrum to dysregulation of behavioural activation: a perspective from dynamical modelling. PLOS One, 8:e63345.

Stetler C, Miller GE (2011). Depression and hypothalamic-pituitary-adrenal activation: a quantitative summary of four decades of research. Biopsychosoc Sci Med, 73:114–126.

Takaesu Y (2018). Circadian rhythm in bipolar disorder: A review of the literature. Psychiatry Clin Neurosci, 72:673-682.

Walker WH, Walton JC, DeVries AC, Nelson RJ (2020). Circadian rhythm disruption and mental health. Transl Psychiatry, 10:28.

